# Bayesian Inference of Other Minds Explains Human Choices in Group Decision Making

**DOI:** 10.1101/419515

**Authors:** Koosha Khalvati, Seongmin A. Park, Saghar Mirbagheri, Remi Philippe, Mariateresa Sestito, Jean-Claude Dreher, Rajesh P. N. Rao

## Abstract

To make decisions in a social context, humans have to predict the behavior of others, an ability that is thought to rely on having a model of other minds known as theory of mind. Such a model becomes especially complex when the number of people one simultaneously interacts is large and the actions are anonymous. Here, we show that in order to make decisions within a large group, humans employ Bayesian inference to model the “mind of the group,” making predictions of others’ decisions while also considering the effects of their own actions on the group as a whole. We present results from a group decision making task known as the Volunteers Dilemma and demonstrate that a Bayesian model based on partially observable Markov decision processes outperforms existing models in quantitatively explaining human behavior. Our results suggest that in group decision making, rather than acting based solely on the rewards received thus far, humans maintain a model of the group and simulate the group’s dynamics into the future in order to choose an action as a member of the group.

## 1 Introduction

The importance of social decision making in human behavior has spawned a large body of research in social neuroscience and decision making (Sanfey, 2007; Camerer, 2011; Henrich et al., 2005; Behrens et al., 2009; Rilling and Sanfey, 2011; Dunne and ODoherty, 2013; Ruff and Fehr, 2014; Joiner et al., 2017). Human behavior relies heavily on predicting future states of the environment under uncertainty and choosing appropriate actions to achieve a goal. In a social context, the degree of uncertainty about the possible outcomes increases dramatically because the behavior of other human beings can be much more difficult to predict than the physics of the environment.

There are two major approaches for action selection in decision making: “model-free” and “model-based” (Dayan and Daw, 2008; Dayan, 2012; Sutton and Barto, 1998; Daw and Dayan, 2014). In the model-free approach, the decision maker performs actions based on the past history of actions and rewards. At each step, the chosen action is the one with the maximum average (or total) reward obtained from previous trials (McAllister, 1991; Mookherjee and Sopher, 1997; Dickinson and Balleine, 2002). In this approach, there is a direct link between choices and past rewards. On the other hand, in model-based decision making, the subject learns a model of the environment, updates the model based on observations and rewards, and chooses actions based on the current state of the model (Daw et al., 2011; Culbreth et al., 2016). As a result, the relationship between rewards received and current choice is indirect. Besides the history of rewards received, the learned model can include other factors such as potential future rewards and more general rules about the environment. Therefore, the model-based approach is more flexible than model-free decision making (Doll et al., 2012; Dolan and Dayan, 2013). However, model-based decision making requires more cognitive resources, for example, for simulation of future events. Thus, there is an inherent trade-off between the two approaches, and determining which approach humans adopt for different situations is an important open area of research.

Several studies have presented evidence in favor of the model-based approach by quantifying the similarity between probabilistic model-based methods and human behavior when the subject interacts with or reasons about another human (Ray et al., 2009; Yoshida et al., 2010; Xiang et al., 2012; Hula et al., 2015; Baker et al., 2017). Additional evidence in favor of a model-based approach comes from studies suggesting that ToM, autobiographical memory, and thinking about the future are all supported by a network of interconnected nodes in the brain known as the Default Mode Network (DMN) (Spreng et al., 2009; Spreng and Grady, 2010).

Compared to reasoning about a single person, decision making in a group with a large number of members can get complicated. On the one hand, having more group members disproportionately increases the cognitive cost of tracking minds (as in model-based approaches) compared to the cost of tracking the reward history of each action (as in model-free approaches). On the other hand, consistent with the importance that human society places on group decisions, a model-based approach might be the optimal strategy.

How does one extend a model-based approach for reasoning about a single person to the case of decision making within a large group? Group decision making becomes even more challenging when the actions of others in the group are anonymous (e.g., voting as part of a jury) (Park et al., 2017). In such situations, reasoning about the state of mind of individual group members is not possible but the dynamics of group decisions do depend on each individual’s actions.

To investigate these complexities that arise in group decision making, we focused on the Volunteer’s Dilemma task, wherein a few individuals endure some costs to benefit the whole group (Diekmann, 1985; Archetti and Scheuring, 2011). Examples of the task include guarding duty, blood donation, and stepping forward to stop an act of violence in a public place (Darley and Latane, 1968). In order to mimic the Volunteer’s Dilemma in a laboratory setting, we use the binary version of a multiround Public Goods Game (PGG) where the actions of each individual are hidden from others (Olson, 1971; Palfrey and Rosenthal, 1984; Diekmann, 1985; Archetti, 2009b,a;Archetti and Scheuring, 2011).

Using an optimal Bayesian framework based on Partially Observable Markov Decision Processes (POMDPs) (Kaelbling et al., 1998), we propose that in group decision making, humans simulate the “mind of the group” by modeling an *average group member* when making their current choices. Our model incorporates prior knowledge, current observations, and a simulation of the future for modeling human decisions within a group. We compared our model to a model-free approach based on the reward history of each action as well as a state-of-the-art descriptive method for fitting human behaviour in the Public Goods Game. Our model explains human behaviour significantly better than the model-free and descriptive approaches. Furthermore, by leveraging the interpretable nature of our model, we are able to show a potential underlying computational mechanism for the group decision making process.

## 2 Results

### 2.1 Human Behavior in Binary Public Goods Game

Participants were 29 adults (mean age 22.97 years old ± 0.37, 14 women). We analyzed the behavioral data of 12 Public Goods Games (PGGs) in which participants played 15 rounds of the game within the same group of *N* players (*N* = 5).

At the beginning of each round, 1 monetary unit (MU) was endowed (E) to each player. In each round, a player could choose between two options: *contribute or* free-ride. Contribution had a cost of C = 1 MU, implying that the player could choose between keeping their initial endowment or giving it up. In contrast to the classical PGG where the group reward is a linear function of total contributions (Fehr and Gachter, 2000; Krajbich et al., 2009; Hauert et al., 2002), in our PGG, public goods were produced as the group reward (G = 2 MU to each player) if and only if at least *k* players each contributed 1 MU. *k* was set to 2 or 4 randomly for each session and conveyed to group members before the start of the session. The resultant reward after one round was E - C + G = 2 MU for the contributor and E + G = 3 MU for the free-rider when public good was produced (the round was a SUCCESS). On the other hand, the contributor had E – C = 0 MU and the free-rider had E = 1 MU when no public good was produced (the round was a FAILURE).

Figure 1 depicts one round of the PGG task. After the subject acted, the total number of contributions, free rides, and the overall outcome of the round was revealed (success or failure in securing the 2 MU group reward) but each individual players actions remained unknown. Although subjects were told that they were playing with other humans, in reality, they were playing with a computer that generated the actions of all the other *N* − 1 = 4 players based on an algorithm (see Methods). In each session, the subject played with a different group of players.

**Figure 1:**
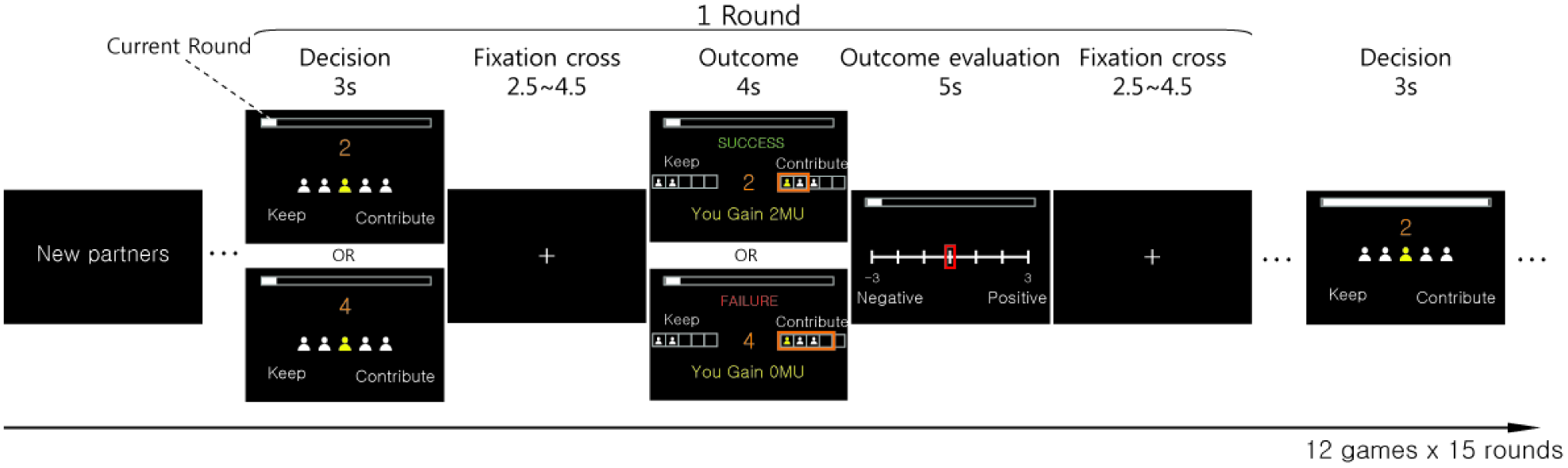
Multi-Round Public Goods Game (PGG). The figure depicts the sequence of computer screens a subject sees in one round of the PGG. The subject is assigned 4 other players as partners and each round requires the subject to make a decision: Keep 1 monetary unit (i.e., free-ride) or contribute. The subject knows whether the threshold to generate public goods (reward of 2 MU for each player) is 2 or 4 contributions (from the 5 players). After the subject acts, the total number of contributions and overall outcome of the round (SUCCESS or FAILURE) are revealed.

As shown in Figure 2a, subjects contributed significantly more when the number of required volunteers was higher with an average contribution rate of 55% (*SD* = .31) for *k* = 4 in comparison to 33% (*SD* = .18) for *k* = 2 (one-tailed paired sample t-test, *t* = 3.94, *df* = 29, *p* = 0.00025, 95% CI difference = [0.08, 0.36]). In addition, Figure 2b shows that the probability of generating public good was significantly higher when *k* = 2 with a SUCCESS rate of 87% (*SD* = 0.09) compared to 36% (*SD* = .29) when *k* = 4 (one-tailed paired sample t-test, *t* = 10.08, *df* = 29, *p* = 4.0 × 10^*−*11^, 95% CI difference =[0.39, 0.62]) (Figure 2b). All but 6 of the subjects contributed more when *k* = 4 (Figure 2c). Of these 6 players, 5 chose to free-ride more than 95% of the time. Also, SUCCESS rate was higher when *k* = 2 for all players (Figure 2d).

**Figure 2:**
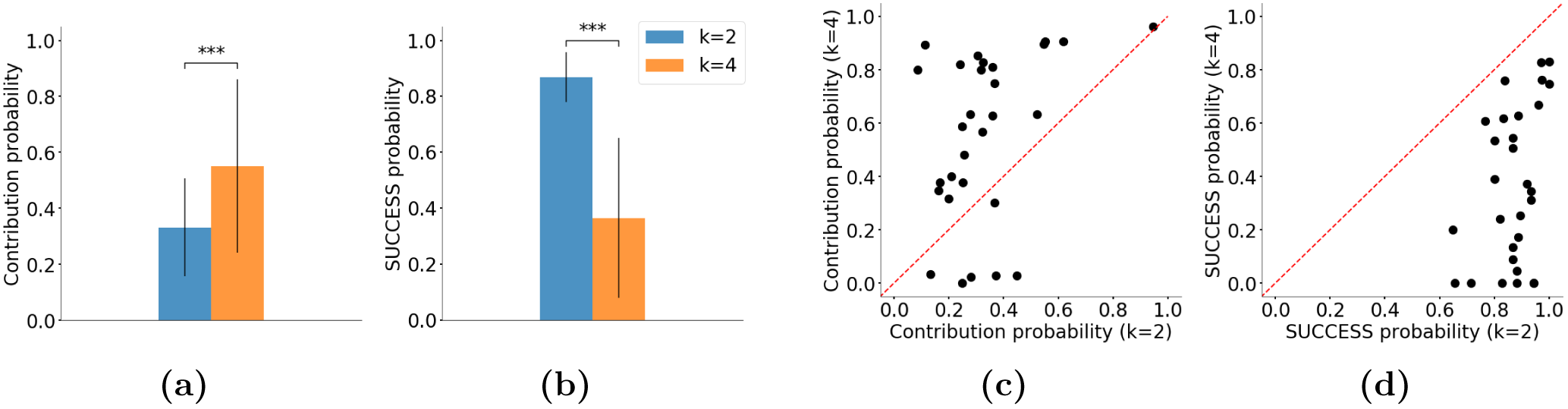
Human Behavior in the PGG Task. (a) The average contribution probability across subjects is significantly higher when the task requires more volunteers (*k*) to generate the group reward. (b) Average probability of SUCCESS across all subjects in generating the group reward is significantly higher when *k* is lower. (c) Average probability of contribution for each subject for *k* = 2 versus *k* = 4. Each point represents a subject. Subjects tend to contribute more often when the task requires more volunteers. (d) Average SUCCESS rate for each subject was higher for *k* = 2 versus *k* = 4.

Average contribution did not change significantly as subjects played more games. In each condition, most of the players played at least 5 games (27 for *k* = 2 and 26 for *k* = 4). For *k* = 2, in their first game, the average contribution rate of players was .37 (*SD* = .25) while in their fifth game, it was .30 (*SD* = .24) (one-tailed paired sample t-test, *t* = 1.34, *df* = 27, *p* = 0.095). When *k* = 4, the average contribution rate was .57 (*SD* = .30) in the first game and .61 (*SD* = .35) in the fifth game (one-tailed paired sample t-test, *t* = *−*0.69, *df* = 26, *p* = 0.25).

### 2.2 Probabilistic Model of Theory of Mind for the Public Goods Game

Consider one round of the PGG task. A player can be expected to choose an action (*contribute* or *free ride*) based on the number of contributions they anticipate the others to make in that round. Since the actions of individual players remain unknown through the game, the only observable parameter is the total number of contributions. One can therefore model this situation using a single random variable *θ*, denoting the average probability of contribution by any group member. With this definition, the total number of contributions by all the other members of the group can be expressed as a binomial distribution. Specifically, if *θ* is the probability of contribution by each group member, the probability of observing *m* contributions from the *N –* 1 others in a group of *N* people is:

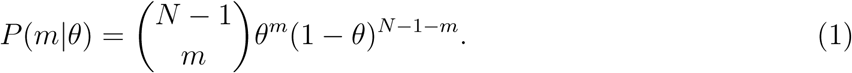

Using this probability, a player can calculate the expected number of contributions from the others, compare it with *k* and decide whether to contribute or free-ride accordingly. For example, if *θ* is very low, there is not a high probability of observing *k* – 1 contributions by the others, implying free riding is the best strategy.

There are two important facts that make this decision making more complex. First, the player does not know *θ*. *θ* must be estimated from the behavior of the group members. We assume that there is a probability distribution over *θ* in the player’s mind, representing their belief about the cooperativeness of the group. Each player starts with an initial probability distribution, called the prior belief about *θ* and updates this belief over successive rounds based on the actions of the others. The prior belief may be based on the previous life experience of the player, or what they believe others would do through fictitious play (Brown, 1951). For example, the player may start with a prior belief that the group will be a cooperative one but change this belief after observing low numbers of contributions by the others. Such belief updates can be performed using Bayes’ rule to invert the probabilistic relationship between *θ* and the number of contributions given by Equation 1.

A suitable prior probability distribution for estimating the parameter *θ* of a Binomial distribution is the Beta distribution, which is itself determined by two (hyper) parameters *α* and *β*:

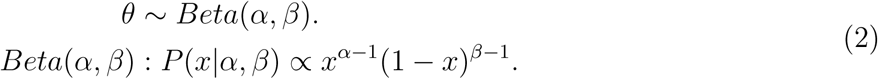

Starting with a prior probability *Beta*(*α*_1_*, β*_1_) for *θ*, the player updates their belief about *θ* after observing the number of contributions from the others in each round through Bayes’ rule. This updated belief is called the posterior probability of *θ*. The posterior probability of *θ* in each round serves as the prior for the next round. With the Beta distribution as the prior and the Binomial distribution as the distribution of observations (here, the number of contributions), the posterior probability distribution remains a Beta distribution (details in Methods) (Murphy, 2012). In our case, with a prior of *Beta*(*α_t_, β_t_*) after observing *c* contributions (including one’s own when made) in round *t*, the posterior probability of *θ* becomes *Beta*(*α_t_*_+1_*, β_t_*_+1_) where *α_t_*_+1_ = *α_t_* +*c* and *β_t_*_+1_ = *β_t_* + *N* – *c*. Note that we include one’s own action in the update of the belief because one’s own action can change the future contribution level of the others.

Intuitively, *α* represents the number of contributions made thus far and *β* the number of free rides. *α*_1_ and *β*_1_ (that define prior belief) represent the player’s a priori expectation about the relative number of contributions versus free-rides respectively before the session begins. For example, when *α*_1_ is larger than *β*_1_, the player starts the task with the belief that people will contribute more than free ride. Large values of *α*_1_ and *β*_1_ imply that the player comes to the game with a lot of prior observations about other players and therefore, the player’s belief will not change significantly after one round of the game when updated with the relatively small number *c* as above.

Decision making in the PGG task is also made complex by the fact that the actual cooperativeness of the group itself (not just the player’s belief about it) may change from one round to the next: players observe the contributions of the others and may change their own strategy for the next round. For example, players may start the game making contributions but change their strategy to free riding if they observe a large number of contributions by the others. We model this phenomenon using a parameter *γ* ≤ 1, which serves as a discount factor for the prior: the prior probability for round *t* is modeled as *Beta*(*γα_t_, γβ_t_*), which allows recent observations about the contributions of other players to be given more importance than observations from the more distant past. Thus, in a round with *c* total contributions (including the subject’s own contribution when made), the subject’s belief about the cooperativeness of the group as a whole changes from *Beta*(*α_t_, β_t_*) to *Beta*(*α_t_*_+1_*, β_t_*_+1_) where *α_t_*_+1_ = *γα_t_* + *c* and *β_t_*_+1_ = *γβ_t_* + *N* – *c*.

### 2.3 Action Selection

How should a player decide whether to contribute or free ride in each round? One possible strategy is to maximize the reward for the current round by calculating the expected number of contributions by the others based on the current belief. Using Equation 1 and the prior probability distribution over *θ*, the probability of seeing *m* contributions by the others when the belief about the cooperativeness of the group is *Beta*(*α, β*) is given by:

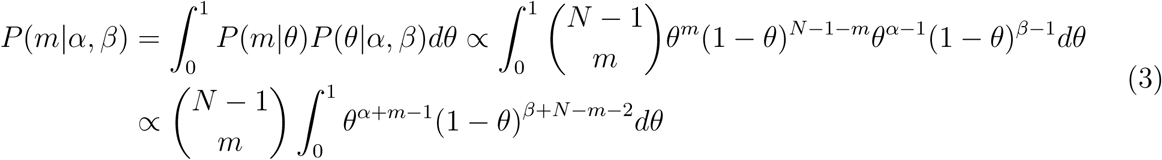

One can calculate the expected reward for the *contribute* versus *free-ride* actions in the current round based on the above equation. Maximizing this reward however is not the best strategy. As alluded to earlier, the actions of each player can change the behavior of other group members in future rounds. The optimal strategy therefore is to calculate the cooperativeness of the group through the end of the session and consider the reward over all future rounds in the session before selecting the current action. Such long-term reward maximization based on probabilistic inference of hidden state in an environment (here, *θ*, the probability of contribution of group members) can be modelled using the framework of Partially Observable Markov Decision Processes (POMDPs) (Kaelbling et al., 1998). Further details can be found in Methods but briefly, to maximize the total expected reward, Markov decision processes start from the last round, calculate the reward for each action and state, and then step back one time step to find the optimal action for each state in that round. This process is repeated in a recursive fashion. Figure 3a shows a schematic of the PGG experiment modeled using a POMDP, while Figure 3b illustrates the mechanism of action selection in this model.

**Figure 3:**
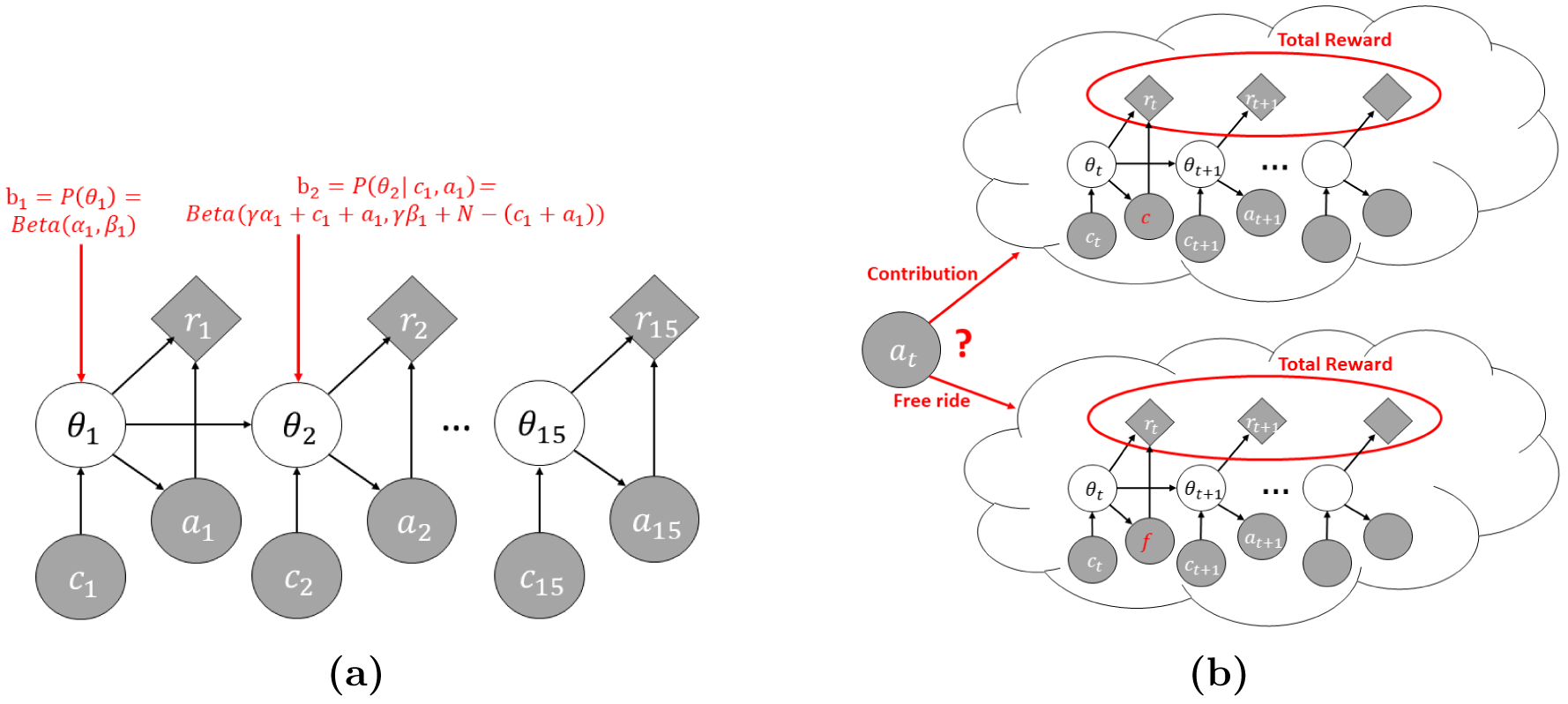
POMDP Model of the Multi-Round Public Goods Game. (a) The subject does not know the average probability of contribution of the group. The POMDP model assumes the subject maintains a probability distribution (”belief”) about the group’s average probability of contribution (denoted here by *θ*) and updates this belief about *θ* after observing the outcome of each round. (b) The POMDP model chooses an action that maximizes total expected reward across all rounds based on the current belief and the consequence of the action on the group behavior in future rounds.

As an example of the POMDP model’s ability to select actions for the PGG task, Figures 4a and 4b show the best actions for a given round (here, round 9) as prescribed by the POMDP model for *k* = 2 and *k* = 4 respectively (the number of minimum volunteers needed); the best actions are shown as a function of different belief states the subject may have, expressed in terms of the different values possible for belief parameters *α_t_* and *β_t_*. This mapping from belief to actions is called a *policy*.

**Figure 4:**
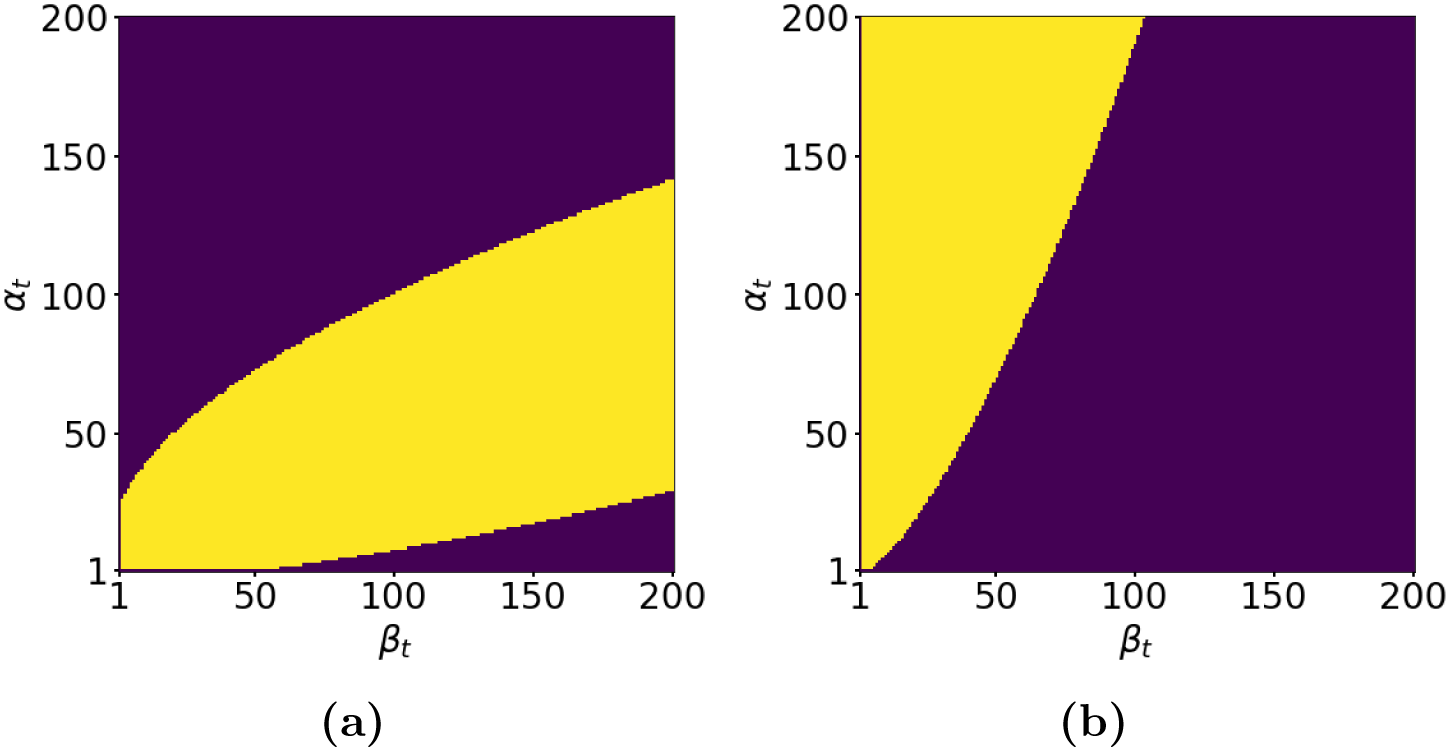
Optimal Actions prescribed by the POMDP Policy as a Function of Belief State. Plot (a) shows the policy for *k* = 2 and plot (b) for *k* = 4. The purple regions represent those belief states (defined by *α_t_* and *β_t_*) for which free-riding is the optimal action; the yellow regions represent belief states for which the optimal action is contributing. These plots confirm that the optimal policy depends highly on *k*, the number of required volunteers. For the two plots, the decay rate was 1 and *t* was 9.

Our simulations using the POMDP model showed that considering a much longer horizon (50 rounds) instead of just 15 rounds gave a much better fit to the subjects’ behavior. Such a long horizon for determining the optimal policy makes the model similar to an infinite horizon POMDP model (Thrun et al., 2005). As a result, the optimal policy for all rounds in our model is very similar to the policy for round 9 shown in Figures 4a and 4b.

### 2.4 POMDP Model Explains Human Behaviour in Volunteer’s Dilemma Task

The POMDP model has three parameters, *α*_1_, *β*_1_, and *γ* which determine the subject’s actions and belief in each round. We fitted these parameters to the subject’s actions by minimizing the error, i.e. the difference between the POMDP model’s predicted action and the subject’s action in each round. The average percentage error across all rounds is then the percentage of rounds that the model predicts incorrectly (*contribute* instead of *free-ride* or vice versa). We defined accuracy as the percentage of the rounds that the model predicts correctly. We also calculated the leave-one-out cross validated (LOOCV) accuracy of our fits (Murphy, 2012).

We found that the POMDP model had an average accuracy across subjects of 84% (*SD* = 0.06) while the average LOOCV accuracy was 77% (*SD* = 0.08). Figure 5a compares the average accuracy and LOOCV accuracy of the POMDP model with two other models. The first is a “model-free” reinforcement learning model known as Q-learning: actions are chosen based on their rewards in previous rounds (Tsitsiklis, 1994) with the utility of group reward, initial values, and learning rate as free parameters (5 parameters per subject – see Methods). The average accuracy of the Q-learning model was 79% (*SD* = .07) which is significantly worse than the POMDP model’s accuracy given above (one-tailed paired t-test, *t* = −6.75, *df* = 29, *p* = 1.26 × 10^*−*7^, 95% CI difference = [0.08, 0.01]). Also, the average LOOCV accuracy of the POMDP model was significantly higher than the average LOOCV accuracy of Q-learning, which was 73% (*SD* = .09) (one-tailed paired t-test, *t* = 2.20, *df* = 29, *p* = 0.0187, 95% CI difference =[0.00, 0.09]).

**Figure 5:**
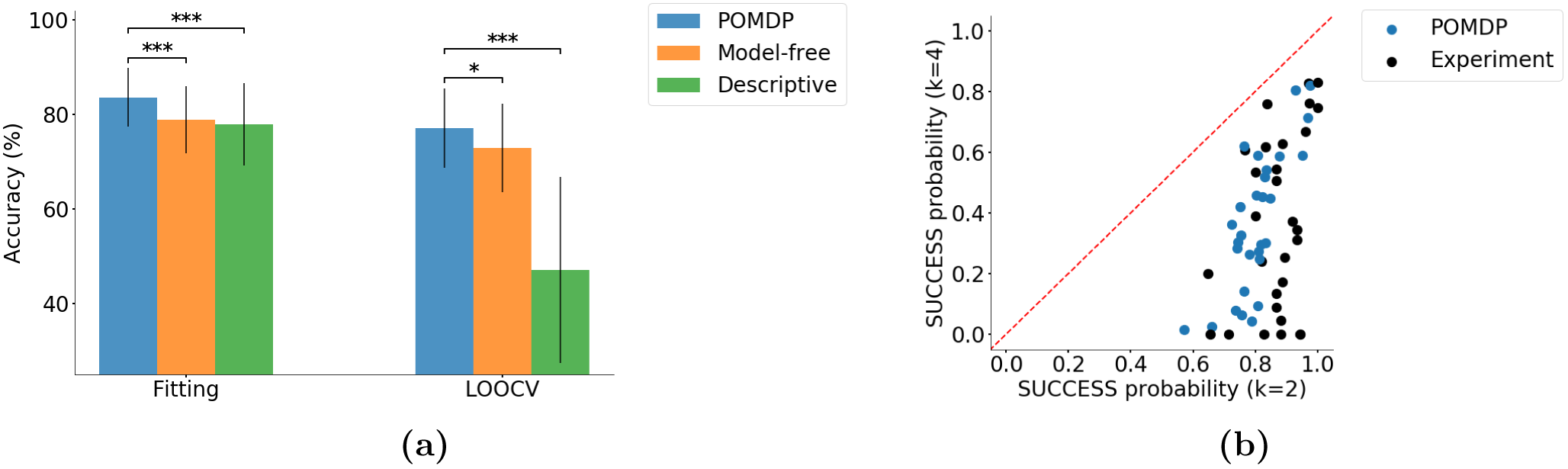
POMDP Model’s Performance and Predictions. (a) Average and LOOCV accuracy across all models. The POMDP model has significantly higher accuracy compared to the other models. (b) POMDP prediction of a subject’s belief about group success in each round (on average) for different *k*’s (blue circles) compared to actual data (black circles, same data as in figure 2d).

We additionally tested a state-of-the-art descriptive model known as the linear two-factor model (Wunder et al., 2013), which predicts the current action of each player based on the player’s own action and contributions by the others in the previous round (this model has three free parameters per subject – see Methods). The average accuracy of the two-factor model was 78% (*SD* = 0.09) which is significantly lower than the POMDP model’s accuracy (one-tailed paired t-test, *t* = −4.86, *df* = 29, *p* = 2.1 × 10^*−*5^, .95% CI difference = [–0.10*, –*0.02]. Moreover, the LOOCV accuracy of the two-factor model was 47% (*SD* = 20), significantly lower than the POMDP model (one-tailed paired t-test, *t* = −7.61, *df* = 29, *p* = 1.4 × 10^*−*8^, 95% CI difference = [–0.38*, –*0.22]). The main reason for this result, especially the lower LOOCV accuracy, is that group SUCCESS also depends on the required number of volunteers (*k*). This value is automatically incorporated in the POMDP’s calculation of expected reward. Also, reinforcement learning works directly with rewards and therefore does not need explicit knowledge of *k* (however, a separate parameter for each *k* is needed in the initial value function for Q-learning – see Methods).

The POMDP model, when fit to a subject’s actions, can also explain other events during the PGG in contrast to the other models described above. For example, based on equation 3 and the action chosen by the POMDP model, one can predict the subject’s belief about the probability of SUCCESS in the current round. This prediction cannot be directly validated but it can be compared to actual SUCCESS. If we consider actual SUCCESS as the ground truth, the average accuracy of POMDP model’s prediction across subjects was 71% (*SD* = .07). Moreover, the prediction matched the pattern of SUCCESS rate data from the experiments (Figure 5b). The other models presented above are not capable of making such a prediction.

### 2.5 Distribution of POMDP Parameters

We can gain insights into the subject’s behavior by interpreting the parameters of our POMDP model in the context of the task. As alluded to above, the prior parameters *α*_1_ and *β*_1_ represent the subject’s prior expectations of contributions and free-rides respectively. Therefore, the ratio *α*_1_*/β*_1_ characterizes the subject’s expectation of contributions by group members while the average of these parameters, (*α*_1_ + *β*_1_)*/*2, indicates the weight the subject gives to prior experience with similar groups before the start of the game. The discount factor (or decay rate) *γ* determines the weight given to past observations compared to new ones: the smaller the discount factor, the more weight the subject gives to new observations. We examined the distribution of these parameter values for our subjects after fitting the POMDP model to their behavior. The ratio *α*_1_*/β*_1_ was in the reasonable range of .5 to 2 for almost all subjects (Figure 6a; in theory the ratio can be as high as 200 or as low as 1*/*200 – see Methods). The value of (*α*_1_ + *β*_1_)*/*2 across subjects was mostly between 40 to 120 (Figure 6b) suggesting that prior belief about groups did have a significant role in players’ strategy, but it was not the only factor since observations over multiple rounds can still alter this initial belief. We also calculated the expected value of contribution by the others in the first round, which is between 0 and *N* – 1 = 4 based on the values of *α*_1_ and *β*_1_ for the subjects. For almost all subjects, this expected value was between 2 and 3 (Figure 6c). Moreover, the discount factor *γ* was almost always above .95, with a mean of .93 and a median of .97 (Figure 6d). Only three subjects had a discount factor less than .95 (not shown in the figure), suggesting that almost all subjects relied on observations made across multiple rounds when computing their beliefs rather than reasoning based solely on the current or most recent observations.

**Figure 6:**
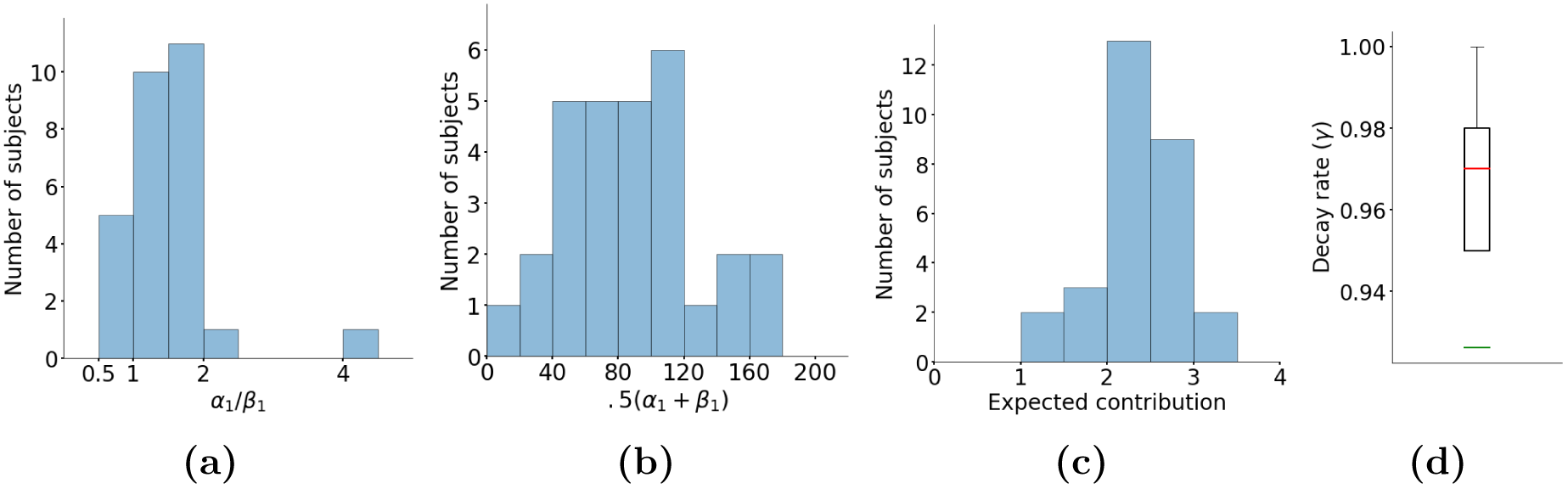
Distribution of POMDP Parameters across Subjects. (a) Histogram of ratio *α*_1_*/β*_1_ shows a value between .5 and 2 for almost all subjects. (b) Histogram of average of *α*_1_ and *β*_1_. For the majority of subjects, this value is between 40 to 120. (c) Histogram of prior belief *Beta*(*α*_1_*, β*_1_) translated into expected contribution by the others in the first round. Note that the values, after fitting to the subjects’ behavior, are mostly between 2 and 3. (d) The box plot of discount factor or decay rate *γ* across subjects shows that this value is almost always above .95. The median is .97 (orange line) and the mean is .93 (green line).

We also calculated each subject’s prior belief about group SUCCESS (probability of SUCCESS in the first round) based on *α*_1_, *β*_1_, and the subject’s POMDP policy in the first round. As group SUCCESS depends on the required number of volunteers (*k*), probability of SUCCESS is different for *k* = 2 and *k* = 4 even with the same *α*_1_ and *β*_1_. Figures 7a and 7b show the distribution of this prior probability of SUCCESS across all subjects for *k* = 2 and *k* = 4. For *k* = 2, all subjects expected a high probability of SUCCESS in the first round, whereas a majority of the subjects expected less than 60% chance for SUCCESS when *k* = 4. While these beliefs cannot be directly validated, the results point to the importance of the required number of volunteers in shaping the subjects’ behavior.

**Figure 7:**
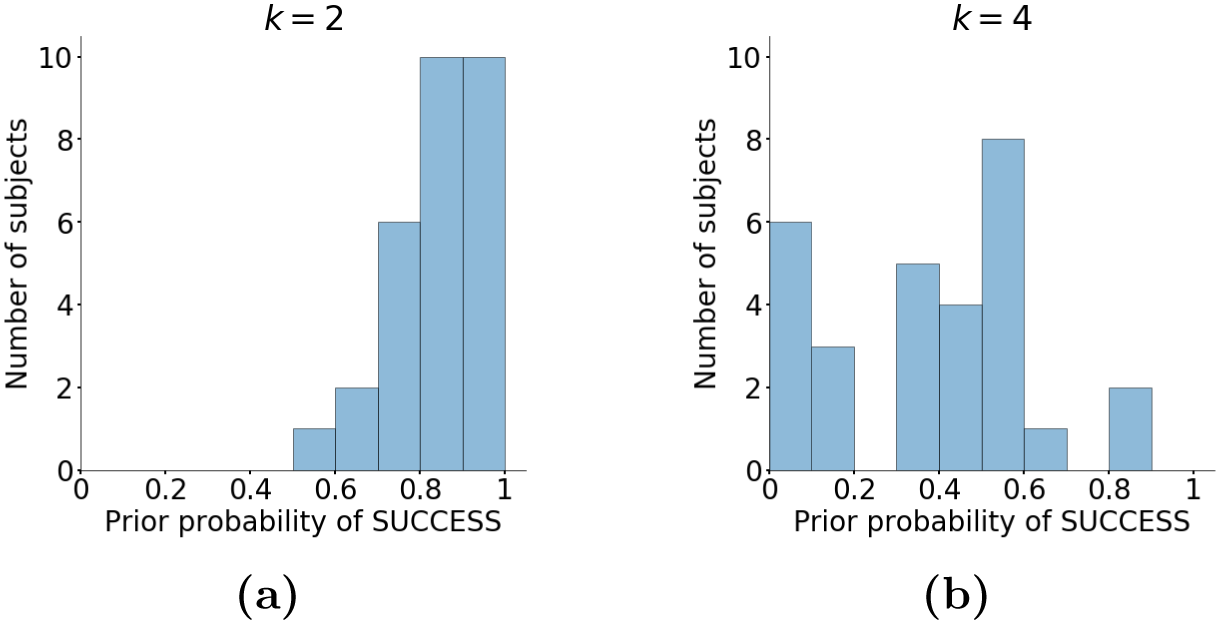
Prior Belief about the Group SUCCESS. (a) When *k* = 2, all subjects expected a high probability of group SUCCESS in the first round (before making any observations about the group). (b) When *k* = 4, almost all subjects assigned a chance of less than 60% to group SUCCESS in the first round.

## 3 Discussion

We introduced a normative model based on POMDPs for explaining human behavior in a group decision making task. Our model combines probabilistic reasoning about the group with long-term reward maximization by simulating the effect of each action on the future behaviour of the group. The greater accuracy of our model in explaining the subjects’ behaviour compared to the other models suggests that humans make decisions in group settings by reasoning about the group as a whole. This mechanism is analogous to maintaining a theory of mind about another person, except theory of mind pertains to a group member on average.

This is the first time, to our knowledge, that a normative model has been proposed for a group decision making task. All existing models to explain human behaviour in the Public Goods game, for example, are descriptive and do not provide insights into the computational mechanisms underlying the decisions (Wanga et al., 2012; Wunder et al., 2013; Fischbacher et al., 2001). Moreover, these methods are based on linear regression of previous events to predict the subject’s action in the current round. These regression-based methods assume independent data points but in a multi-round game, the rounds are not independent. Nonetheless, our model outperforms this family of methods as well.

In addition to providing a better fit to the subject’s behavior, our model, when fit to the subject’s actions can predict SUCCESS rate in each round without being explicitly trained for such predictions, in contrast to the other methods. Also, as alluded to in figures 6a, 6b and 6d, when fit to the subjects’ actions, the parameters were all within a reasonable range, showing the importance of prior knowledge and multiple observations in decision making. These observations imply that our high accuracy in explaining subjects’ behaviour is not an artifact of the complexity of the model or of the Beta distribution.

The POMDP policy aligns with our intuition about action selection in the Volunteer’s Dilemma task. A player chooses to free ride for two reasons: (i) when the cooperativeness of the group is low and therefore there is no benefit in contributing, and (ii) when the player knows there are already enough volunteers and contributing leads to a waste of resources. The two purple areas of Figure 4a represent these two conditions for *k* = 2. The upper left part represents large *α_t_* and small *β_t_*, implying a high contribution rate, while the bottom right part represents small *α_t_* and large *β_t_* implying a low contribution rate. When *k* = 4, all but one of the 5 players must contribute for group SUCCESS - this causes a significant difference in the optimal POMDP policy compared to the *k* = 2 condition. As seen in Figure 4b, there is only a single region of belief space for which free-riding is the best strategy, namely, when the player does not expect contributions by enough players (relatively large *β_t_*). On the other hand, as expected, this region is much larger compared to the same region for *k* = 2 (see Figure 4a). The POMDP model predicts that free-riding is not a viable action in the *k* = 4 case (Figure 4b) because not only does this action require all the other 4 players to contribute in order to generate the group reward in the current round, but such an action also increases the chances that the group contribution will be lower in the next round, resulting in lesser expected reward in future rounds.

In a game with a predetermined and known number of rounds, even if the player considers the future, one might expect the most rewarding action in the last rounds to be free riding as there is little or no future to consider. However, our data did not support this conclusion. Our model is able to explain the data by using a longer horizon than the number of rounds in the game. While this may make the policy deviate from optimality, it provides a significant computational benefit by making the policies for different rounds similar to each other, avoiding re-calculation of a policy for each single round. Recent studies in human decision making have demonstrated that humans may use such minimal modifications of model-based policies for efficiency (Momennejad et al., 2017; Russek et al., 2017). An additional reason behind favoring contribution in the PGG may be an incomplete understanding of the task by some subjects (Burton-Chellew and West, 2013).

Our model assumes that the subject estimates the cooperativeness of others in each round before choosing the next action. It is possible to extend this reasoning by considering the effect of the chosen action on other minds in terms of their cooperativeness, and so on in a recursive fashion. If such a multi-level theory of mind is extended to infinite depth, the game converges to the Nash equilibrium (Nash, 1950). In reality, however, such an infinite-depth theory of mind appears not to occur in actual social interactions among humans (Kagel and Roth, 2016; Camerer, 2011; Henrich et al., 2005), with multi-level theory of mind limited to very few levels as observed in some experiments (Camerer et al., 2004; Yoshida et al., 2008, 2010; Hula et al., 2015). Moreover, we expect less levels of theory of mind in group decision making in general compared to two-person interactions due to the complexity of the group decision making process and the anonymity of individuals.

In the Volunteer’s Dilemma, not only is the common goal not reached when there are not enough volunteers, but having more than the required number of volunteers leads to a waste of resources. As a result, an accurate prediction of others’ intentions based on one’s beliefs is crucial to make accurate decisions. This gives the model-based approach a huge advantage over model-free methods in terms of reward gathering, thus making it more beneficial for the brain to endure the extra cognitive cost. It is possible that in simpler tasks where the accurate prediction of minds is less crucial, the brain adopts a model-free approach.

Our model was based on the binomial and beta distributions for binary values due to nature of the task, but it can be easily extended to the more general case of a discrete set of actions using multinomial and Dirichlet distributions. Additionally, the model can be extended to multivariate states, e.g., when the players are no longer anonymous. In such cases, the belief can be modeled a joint probability distribution over all parameters of the state. This however incurs a significant computational cost. An interesting area for future research is investigating whether under some circumstances, humans model group members with similar behaviour as one subgroup in order to reduce the number of minds one should reason about.

A strength of the POMDP model is that it can model social tasks beyond economic decision making, such as prediction of others’ intentions and actions in everyday sitations (Koster-Hale and Saxe, 2013; Tamir and Thornton, 2018). In these cases, we would need to modify the model’s definition of the state of other minds to include dimensions such as valence, competence, and social impact instead of propensity to contribute monetary units as in the PGG task (Posner et al., 2005; Gray et al., 2007; Cuddy et al., 2008; Tamir et al., 2016).

The interpretability of the POMDP framework offers an opportunity to study the neurocognitive mechanisms of group decision making in healthy and diseased brains. POMDPs and similar Bayesian models have previously proved useful in understanding neural responses in sensory decision making (Rao, 2010; Huang et al., 2012; Huang and Rao, 2013; Khalvati and Rao, 2015) and in tasks involving interactions with a single individual (Xiang et al., 2012; Yoshida et al., 2010; Hula et al., 2015; Baker et al., 2017). We believe the POMDP model we have proposed can likewise prove useful in interpreting neural responses and data from neuroimaging studies of group decision making tasks. Additionally, the model can be used for Bayesian theory-driven investigations in the field of computational psychiatry (Huys et al., 2016). For example, ToM deficits are a key feature of autism spectrum disorder (Baron-Cohen et al., 2001) but it is unclear what computational components are impaired and how they are affected. The POMDP model may provide a new avenue for computational studies of such neuropsychiatric disorders.

## 4 Methods

### 4.1 Experiment

30 right-handed students at the University of Parma were recruited for this study. One of them aborted the experiment due to anxiety. Data from the other 29 participants were collected, analyzed, and reported. Based on self-reported questionnaires, none of the participants had a history of neurological or psychiatric disorders. This study was approved by the Institutional Review Board of the local ethics committee from Parma University (IRB no. A13-37030), which was carried out according to the ethical standards of the 2013 Declaration of Helsinki. All participants gave their informed written consent. As mentioned in Results, each subject played 14 sessions of the Public Goods Game (PGG) (i.e., the Volunteer’s Dilemma), each containing 15 rounds. In the first 2 sessions, subjects received no feedback about the result of each round. However, in the following 12 sessions, social and monetary feedback were provided to the subject. The feedback included the number of contributors and free riders, and the subject’s reward in that round. Each individual player’s action, however, remained unknown to the others. Therefore, individual players could not be tracked. We present analyses from the games with feedback.

In each round (see Figure 1), the participant had to make a decision within three seconds by pressing a key; otherwise the round was repeated. 2.5 to 4 seconds after the action selection, the outcome of the round was shown to the subject for 4 seconds. Then, players evaluated the outcome of the round before the next round started. Subjects were told that they were playing with 19 other participants located in other rooms. Overall, 20 players were playing the PGG in 4 different groups simultaneously. These groups were randomly chosen by a computer at the beginning of each session. In reality, subjects were playing with a computer. In other words, a computer algorithm was generating all the actions of others for each subject. Each subject got a final monetary reward equal to the result of one PGG randomly selected by the computer at the end of the study.

In a PGG with *N* = 5 players, we denote the action of player *i* in round *t* with the binary value of 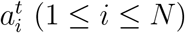 with 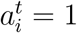 representing contribution and 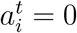 representing free-riding. The human subject is assumed to be player 1. We define the average contribution rate of others 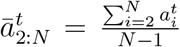 and generate each of the *N* − 1 actions of others in round *t* using the following probabilistic function:

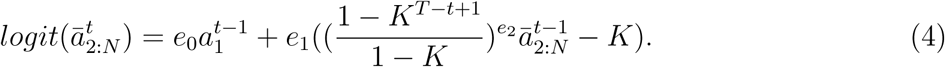

where *K* = *k/N* where *k* is the required number of contributors.

This model has 3 free parameters: *e*_0_*, e*_1_*, e*_2_. These were obtained by fitting the above function to the actual actions of subjects in another PGG study (Park et al., 2013), making this function a simulation of human behavior in the PGG task. For the first round, we used the mean contribution rate of each subject as their fellow members’ decision.

### 4.2 Markov Decision Processes

A Markov Decision Process (MDP) is a tuple (*S, A, T, R*) where *S* represents the set of states of the environment, *A* is the set of actions, *T* is the transition function *S* × *S* × *A* → [0, 1] that determines the probability of the next state given the current state and action, i.e. *T* (*sä, s, a*) = *P* (*sä|s, a*), and *R* is the reward function *S* × *A* → *ℝ* representing the reward associated with each state and action (Bellman, 1957). In an MDP with horizon *H* (total number of performed actions), given the initial state *s*_1_, the goal is to choose a sequence of actions that maximizes the total expected reward:

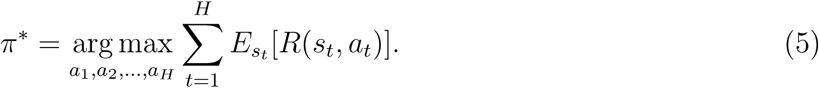

This sequence, called the optimal policy, can be found using the technique of dynamic programming (Thrun et al., 2005). For an MDP with time horizon *H*, define the value function *V* and action function *U* at the last time step *t* = *H* as:

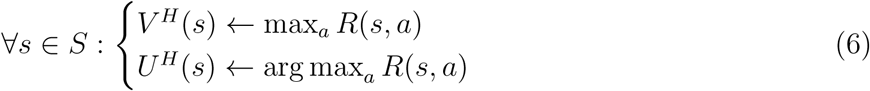

For any *t* from 1 to *H* − 1, the value function *V^t^* and action function *U^t^* is defined recursively as:

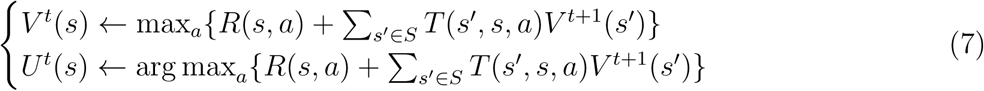

Starting from the initial state *s*_1_ at time 1, the action chosen by the optimal policy *π\** at time step *t* is *U^t^* (*s_t_*).

When the state of the environment is hidden, the MDP turns into a Partially Observable MDP (POMDP) where the state is estimated probabilistically from observations or measurements from sensors. Formally, a POMDP is defined as (*S, A, Z, T, O, R*) where *S, A, T, R* are defined as in the case of MDPs, *Z* is the set of possible observations, and *O* is the observation function *Z* ×*S* → [0, 1] that determines the probability of any observation *z* given a state *s*, i.e., *O*(*z, s*) = *P* (*z|s*). In order to find the optimal policy, the POMDP model uses the posterior probability of states, known as the belief state, where *b_t_*(*s*) = *P* (*s z*_1_*, a*_1_*, z*_2_*, …, a_t−_*_1_). Belief states can be computed recursively as follows:

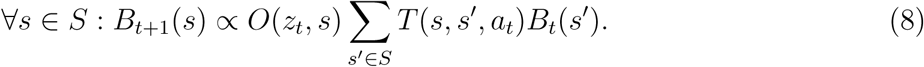

If we define *R*(*b_t_, a_t_*) as the expected reward of *a_t_*, i.e., 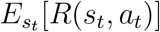, starting from initial belief state, *b*_1_, the optimal policy for the POMDP is given by:

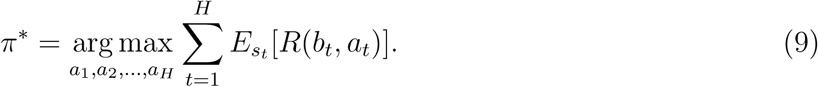

A POMDP can be considered an MDP whose states are belief states. This belief state space however is exponentially larger than the underlying state space. Therefore, solving a POMDP optimally is computationally expensive (Smallwood and Sondik, 1973), unless the belief state can be represented by a few parameters as in our case Thrun et al. (2005). For solving larger POMDP problems, various approximation and learning algorithms have been proposed. We refer the reader to the growing literature on this topic (Ross et al., 2008; Silver and Veness, 2010; Khalvati and Mackworth, 2013; Shani et al., 2013; Luo et al., 2018).

### 4.3 POMDP for Binary Public Goods Game

The state of the environment is represented by the average cooperativeness of the group, or equivalently, the average probability *θ* of contribution by a group member. Since *θ* is not observable, the task is a POMDP and one must maintain a probability distribution (belief) over *θ*. The Beta distribution, represented by two free parameters (*α* and *β*), is the conjugate prior for binomial distribution Murphy (2012). Therefore, when performing Bayesian inference to obtain the belief state over *θ*, combining the Beta distribution as the prior belief and the binomial distribution as the likelihood results in another Beta distribution as the posterior belief. Using the Beta distribution for the belief state, our POMDP turns into an MDP with a two-dimensional state space represented by *α* and *β*. Starting from an initial belief state *Beta*(*α*_1_*, β*_1_) and with an additional free parameter *γ*, the next belief states is determined by the actions of all players at each round as described in Results. For the reward function, we used the monetary reward function of the Public Goods Game (PGG). Therefore, the elements of our new MDP derived from the PGG POMDP are as following:

- *S* = (*α*, *β*)
- *A* = (*c*, *f*)
- 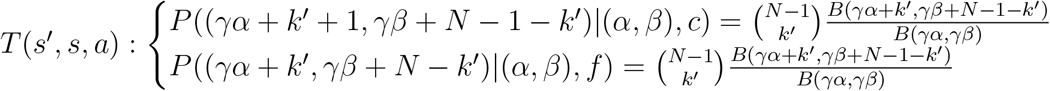
- 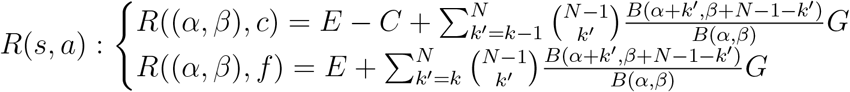
- *B*(*α, β*) is the normalizing constant: 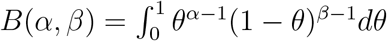.

According to the experiment, the time horizon should be 15 time steps. However, we found that a very long horizon (*H* = 50) for all players provides a better fit to the subjects’ data. For each subject, we found *α*_1_*, β*_1_, and *γ* that made our POMDP’s optimal policy fit the subject’s actions as much as possible. For simplicity, we only considered integer values for states (integer *α* and *β*). The fitting process involved searching over integer values from 1 to 200 for *α*_1_ and *β*_1_ and values between 0 to 1 with a precision of .01 (.01*, .*02*, …, .*99, 1.0) for *γ*. The fitting criterion was round-by-round accuracy. For consistency with the descriptive model, the first round was not included (despite the POMDP model’s capability for predicting it). Since the utility value for public good for a subject can be higher than the monetary reward due to social or cultural reasons (Fehr et al., 2002; Rilling et al., 2002), we investigated the effect of higher values for the group reward *G* in the reward function of the POMDP. This however did not improve the fit.

In round *t*, if the POMDP model selects the action “contribution”, the probability of SUCCESS can be calculated as 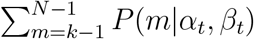 (see Equation 3). Otherwise, the probability of SUCCESS is 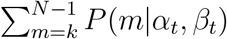. This probability value was compared to the actual SUCCESS and FAILURE of each round to compute the accuracy of SUCCESS prediction by the POMDP model.

A preliminary version of the above model but without the *γ* parameter was presented in (Khalvati et al., 2016).

### 4.4 Model-Free Method: Q-Learning

We used Q-learning as our model-free approach. There are two Q values in the PGG task, one for each action, i.e., Q(c) and Q(f) for “contribute” and “free-ride” respectively. At the beginning of each PGG, Q(c) and Q(f) are initialized to the expected reward for a subject for that action based on a free parameter *p* which represents the prior probability of group SUCCESS. As a result, we have:

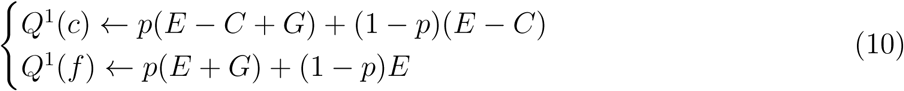

We customized the utility function for each subject by making the group reward *G* a free parameter. Moreover, as the probability of SUCCESS is different for *k* = 2 and *k* = 4, we used two separate parameters *p*_2_ and *p*_4_ instead of *p*, depending on the value of *k* in the PGG.

In each round of the game, the action with the maximum Q value was chosen. The Q value for that action was then updated based on the subject’s action and group SUCCESS/FAILURE, with a learning rate *η^t^*. This learning rate was a function of the round number, i.e. 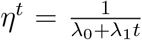 where *λ*_0_ and *λ*_1_ are free parameters and *t* is the number of the current round. Let the subject’s action in round *t* be *a^t^*, the Q-learning model’s chosen action be *â*^*t*^, and the reward obtained be *r^t^*. We have:

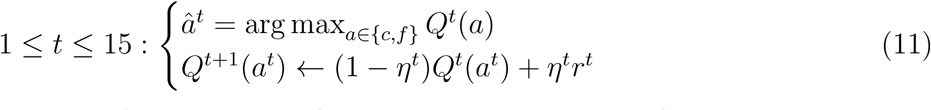

For each subject, we searched for the values of *λ*_0_, *λ*_1_, the group reward *G*, and the probability of group SUCCESS *p*_2_ or *p*_4_ that maximize the round-by-round accuracy of the Q-learning model. Similar to the other models, the first round was not included in this fitting process.

### 4.5 Descriptive Model

Our descriptive model was based on a logistic regression (implementation by Scikit-learn (Pedregosa et al., 2011)) that predicts the subject’s action in the current round based on their own previous action and the total number of contributions by the others in the previous round. As a result, this model has 3 free parameters (two features and a bias parameter). Let 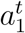 be the subject’s action in round *t* and 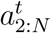 be the actions of others in the same round. The subject’s predicted action in the next round *t* + 1 is then given by:

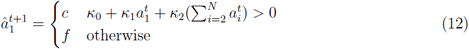

We used one separate regression model for each subject. As the model’s predicted action is based on the previous round’s actions, the subject’s action in the first round cannot be predicted by this model.

### 4.6 Greedy Strategy

If a player wants to solely maximize the expected reward in the current round and ignores the future, the optimal action is always free-riding independent of the average probability of contribution by a group member. This is because free-riding always results in one unit more monetary reward (3 MU for SUCCESS or 1 MU for FAILURE) compared to contribution (2 MU or 0 MU), except in the case where the total number of contributions by others is *exactly k –* 1. In the latter case, choosing contribution yields 1 unit more reward (2 MU) compared to free-riding (1 MU). This means that the expected reward for free-riding is always more than that for contribution unless the probability of observing exactly *k* − 1 contributions by others is greater than .5. We show that this is impossible for any value of *θ*. First, note that the probability of exactly *k* – 1 contributions from *N −* 1 players is maximized when *θ* = (*k* − 1)*/*(*N* − 1). Next, for any *θ*, the probability of *k* − 1 contributions from *N* − 1 players is:

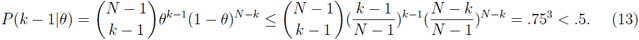

for *N* = 5 and for either *k* = 2 or *k* = 4.

## 5 Acknowledgments

This work was funded by the NSF-ANR Collaborative Research in Computational Neuroscience ‘CRCNS SOCIAL POMDP’ *n°*16-NEUC to J.C.D. and CRCNS NIMH grant no. 5R01MH112166-03, NSF grant no. EEC-1028725, and a Templeton World Charity Foundation grant to R.P.N.R. The experiments were performed within the framework of the Laboratory of Excellence ‘LABEX ANR-11-LABEX-0042’ of Universite de Lyon, attributed to J.C.D., within the program “Investisse-ments d’Avenir” (ANR-11-IDEX-0007) operated by the French National Research Agency (ANR). J.C.D. was also funded by the IDEX University Lyon 1 (project INDEPTH).

## 6 Contributions

R.P.N.R. and J.C.D. developed the general research concept. S.A.P. designed and programmed the task under J.C.D.’s supervision and M.S. ran the experiment under J.C.D.’s supervision. K.K. developed the model under R.P.N.R.’s supervision, implemented the algorithms and analyzed the data in collaboration with R.P.N.R. S.M. interpreted the computational results in the context of social neuroscience. K.K. developed the reinforcement learning model after discussions with R.P. K.K., S.M., and R.P.N.R. wrote the manuscript in collaboration with S.A.P., R.P., and J.C.D.

